# Detection of cerebral tauopathy in P301L mice using high-resolution large-field multifocal illumination fluorescence microscopy

**DOI:** 10.1101/2020.05.18.101188

**Authors:** Ruiqing Ni, Zhenyue Chen, Juan A. Gerez, Gloria Shi, Quanyu Zhou, Roland Riek, K. Peter R. Nilsson, Daniel Razansky, Jan Klohs

## Abstract

Current intravital microscopy techniques visualize tauopathy with high-resolution, but have a small field-of-view and depth-of-focus. Herein, we report a transcranial detection of tauopathy over the entire cortex of P301L tauopathy mice using large-field multifocal illumination (LMI) fluorescence microscopy technique and luminescent conjugated oligothiophenes. *In vitro* assays revealed that fluorescent ligand h-FTAA is optimal for *in vivo* tau imaging, which was confirmed by observing elevated probe retention in the cortex of P301L mice compared to non-transgenic littermates. Immunohistochemical staining further verified the specificity of h-FTAA to detect tauopathy in P301L mice. The new imaging platform can be leveraged in pre-clinical mechanistic studies of tau spreading and clearance as well as longitudinal monitoring of tau targeting therapeutics.

## 1. Introduction

The abnormal deposition of pathological tau fibrils is a characteristic feature of tauopathy related neurodegenerative diseases including Alzheimer’s disease, frontotemporal lobar dementia (FTLD), chronic traumatic encephalopathy, corticobasal degeneration, progressive supranuclear palsy and parkinsonism linked to chromosome 17 [1]. Microtubule associated protein Tau locates intracellularly and is composed of six isoforms classified into 4-repeat (4R) and 3-repeat (3R) species, and the composition of tau isoforms differ among diverse tauopathies [2]. Tau abnormal accumulation in patients with Alzheimer’s disease was found to be closely related to axonal damage, neurodegeneration and cognitive impairment [3–5]. This designates tauopathy an important target for early diagnostic and therapeutic intervention for Alzheimer’s disease, FTLD, and other tauopathy disorders [6]. Tau tracers for positron emission tomography (PET) have been developed recently [5, 7–12], while tau imaging has emerged as a useful tool for disease staging, progression prediction, treatment stratification, and monitoring in clinical research setting.

Several tau mouse models that recapitulate pathological features of tauopathy have been developed with mutations in the *MAPT* gene, including P301S [13], P301L (rTg4510 under CaMKII [14], prion [15] and Thy1.2 [16] promotors), and hTau lines [17]. Whole brain high-resolution tau imaging in these disease models provide insights for mechanistic and therapeutic studies [18, 19]. At macroscopic level, *in vivo* imaging of tauopathy was enabled by PET in P301L mouse line using ^11^C-PBB3 [20–22] or ^18^F-PI-2620 [10] tau probes. However, the limited resolution of microPET (1 mm) relative to the mouse brain (~10 mm^3^) and the need for dedicated and expensive infrastructure for radiolabeling limit usability in preclinical setting. Near-infrared fluorescence imaging using PBB5, PBB3 [20], fluorescently labelled antibodies, and antibody fragments [23] have also been applied for tauopathy detection in animal models. Deep tissue fluorescence imaging is yet affected by severe light scattering and absorption, resulting in limited imaging depth as well as poor spatial resolution and quantification accuracy. At the microscopic scale, two-photon imaging with cranial window using Congo-red derivative FSB [24, 25], Thioflavin S [26], and luminescent conjugated oligothiophene (LCO) HS-84 [27] have monitored tauopathy in mouse models with cellular/sub-cellular resolution but restricted (sub-millimeter) field-of-view (FOV).

The aim of the present study is to assess the selectivity and specificity of LCOs [28] for *in vivo* tauopathy detection in the P301L (Thy1.2) mouse model [16] with large-field multifocal illumination (LMI) fluorescence microscopy attaining 12×12 mm FOV and high spatial resolution (~6 μm) [29, 30].

## 2. Materials and methods

### 2.1 LMI fluorescence imaging system and characterization

The LMI fluorescence imaging system is based on a beam-splitting grating and an acousto-optic deflector, which are synchronized with a high speed camera to attain real-time fluorescence microscopy over a 20×20 mm FOV [29, 30]. A 473 nm CW laser was used for h-FTAA excitation. The laser beam was first scanned by a two dimensional acousto-optic deflector (AA Opto-Electronic, France) at 900 Hz and then guided into a customized beam-splitting grating (Holoeye GmbH, Germany) to generate 21×21 mini-beam pattern. It is subsequently focused with a scan lens (CLS-SL, Thorlabs) to generate the illumination grid upon the sample. After passing through the dichroic mirror, the emitted fluorescence signals are collected by a Nikon lens (Micro-Nikkor AF-S 60mm f/2.8 G, Nikon) and focused onto the sensor plane of a high-speed camera (pco.dimax S1, PCO AG, Germany). To render one LMI image, the illumination grid is raster scanned over 90×90 positions with ~7 μm scanning step. After data acquisition, only signals at the excitation foci were extracted and then superimposed to form the high-resolution image.

### 2.2 In vitro binding of probes to recombinant tau fibrils and mouse brain

Recombinant amyloid-beta Aβ_42_ and full-length tau were expressed and produced by *E.coli* as described previously [31, 32]. The fluorescence LCO probes h-FTAA, q-FTAA, HS-84 and HS-169 were synthesized as described previously [33, 34]. Thioflavin T assays using LCOs (q-FTAA, q-FTAA, HS-84 and HS-169) at 1μM, AOI987 at 10 nM against Aβ_42_ (80.3 μg/ml) and tau fibrils (379.2 μg/ml) using fluorometer (Fluoromax 4, Horiba scientific, Japan) were performed as described previously [30], with three technical replicates. These experiments were conducted twice.

### 2.3 Animal model

Two 17 months-old and four 10 months-old mice transgenic for MAPT P301L, overexpressing human 2N/4R tau under neuron-specific Thy1.2 promoter (pR5 line, C57B6.Dg background) [16, 35], and four non-transgenic littermates were used (4 and 10 months-of-age), of both gender. Animals were housed in individually ventilated cages inside a temperature-controlled room, under a 12-hour dark/light cycle. Pelleted food (3437PXL15, CARGILL) and water were provided ad-libitum. All experiments were performed in accordance with the Swiss Federal Act on Animal Protection and were approved by the Cantonal Veterinary Office Zurich (permit number: ZH082/18).

### 2.4 In vivo and ex vivo LMI fluorescence imaging of tauopathy in P301L mice

For *in vivo* LMI imaging, mice were anesthetized with isoflurane (4 % v/v for induction and 1.5 % v/v during experiments) in O_2_: air (1/4) mixture at a flow rate of ~0.8 l/min. Lidocaine 7 mg/kg and Epinephrine 7 μg/kg were applied subcutaneously. Before scanning, each mouse was positioned onto the imaging stage, with the scalp removed to reduce light scattering while the skull was kept intact. For P301L and non-transgenic littermate mice, an *i.v.* tail-vein injection of 0.6 mg/kg h-FTAA solution in 0.1 M phosphate buffered saline (PBS, pH 7.4) was administered. LMI imaging and widefield imaging were recorded before and after h-FTAA injection with a time interval of 20 min and terminated at 120 min post-injection.

Seven mice were sacrificed under deep anesthesia (ketamine/xylazine/acepromazine maleate, 75/10/2 mg/kg body weight, *i.p.* bolus injection) without prior perfusion. The dissected skull, the whole brain and brain slices of 5-mm thickness, prepared using brain matrix and razor blade, were imaged with LMI and widefield imaging on object holders wrapped with black tape using the aforementioned set-up. To validate the *in*- and *ex vivo* signal, the other two P301L and non-transgenic littermate mice were perfused under ketamine/xylazine/acepromazine maleate anesthesia (75/10/2 mg/kg body weight, *i.p.* bolus injection) with 0.1 M PBS (pH 7.4) and decapitated. The brains were then imaged using LMI imaging, immersion fixed in 4 % paraformaldehyde in 0.1 M PBS (pH 7.4) for 1 day and then stored in 0.1 M PBS (pH 7.4) at 4 °C for immunohistochemistry.

### 2.5 Image reconstruction and data analysis

Reconstruction of the LMI imaging data was performed based on the saved raw data from the CMOS camera. Firstly, local maxima in each scanning frame were identified and the excited fluorescence signals were extracted with their centroids and intensity stored for the subsequent image compounding. Since the illumination grid was well-defined with equal intervals between adjacent spots, this prior information facilitated signal extraction while suppressing noise. Note that the beamsplitting grating was originally designed for the 532 nm wavelength, resulting in a large uniformity error for the zero diffraction order at the 473 nm wavelength. Thus, signal intensity correction was implemented for the central spot before signal superimposition. According to image scanning microscopy theory [36], each scanning image has to be shifted with respect to a common frame before rendering the final high-resolution compounded image. In our case, the extracted signals at the excitation spots were not altered but placed four times further from each other in the final compounded image given the low magnification power and 90×90 raster scanning positions.

For data analysis, fluorescence intensity (F.I.) in five regions-of-interest (ROIs) were drawn for the *ex vivo* experimental results of both LMI and widefield imaging in P301L mice (Fig. 4a, b, red square: ROI, and blue square: background region) for Contrast-to-noise ratio (CNR) analysis according to Eq. (1).

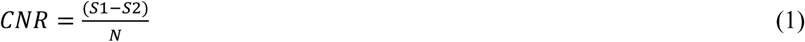

where *S* is F.I. in the ROI; and N is the standard deviation from a region in the background.

### 2.6 Staining and confocal imaging

LCOs were applied to brain tissue slices from one P301L and one non-transgenic littermate. Horizontal brain sections (5 μm) were stained with h-FTAA, q-FTAA, HS-84, HS-169 (5 μM, dd water), and counterstained using DAPI (Sigma, 1:1000 dilution). Confocal images of brain sections from P301L and non-transgenic littermates were obtained using a Leica SP8 confocal microscope (Leica Microsystems GmbH, Germany) at ScopeM ETH Zurich Hönggerberg core facility. Lambda scan using Leica SP8 was performed on h-FTAA and HS-84 stained brain slices to further determine the fluorescent properties of the dyes. Three ROIs were chosen on the tauopathy deposits stained by dyes.

After *ex vivo* LMI imaging, brains from P301L mice and non-transgenic littermate mice were removed from the skull and embedded in paraffin following routine procedures for histology and immunohistochemical investigations. Horizontal and sagittal brain sections (5 μm) were made and stained with 5 μM h-FTAA, and anti-phosphorylated antibodies AT-8 (pSer202/pThr205, MN1020, Invitrogen, 1:50) and AT-100 (pThr212/pSer214, MN1060, Invitrogen, 1:50) [35, 37], conjugated with Alexa647 Goat polyclonal mouse (Invitrogen, A-21235, 1:100). Sections were counterstained using DAPI (1:1000). The brain sections were imaged at ×20 magnification using Panoramic 250 (3D HISTECH, Hungary) and at ×63 magnification using a Leica SP8 confocal microscope facility for co-localization of h-FTAA and AT-8/AT-100. The images were analyzed using CaseViewer (3D HISTECH, Hungary) and ImageJ (NIH, U.S.A).

### 2.7 Statistics

An unpaired two-tail student *t* test was used (Graphpad Prism) for comparing values between two groups. For the time series, two-way ANNOVA was used. All data are present as mean ± standard deviation. Significance was set at \**p* < 0.05.

## 3. Results

### 3.1 In vitro probe characterization

Neurodegenerative diseases are associated not only with the aggregation of tau fibrils, but also with other intra- and extracellular protein aggregates such as Aβ and alpha-synuclein and the ideal probe for tau imaging should be selective for tau fibrils. First, we tested and compared the selectivity of several LCOs and fluorescent probes for their binding to tau fibrils versus Aβ_42_ fibrils. Fluorescence binding assays were performed *in vitro* using a panel of probes binding to Aβ_42_ and full-length (4R/2N) tau fibrils using a fluorescence spectrometer. As expected, Thioflavin-T binds to Aβ42 and tau fibrils, as both proteins have an extensive cross β-pleated sheet conformation [32]. The LCOs h-FTAA, q-FTAA, HS-84, and HS-169 show also binding to Aβ_42_ and tau fibrils. In contrast, AOI987 designed to target Aβ [38, 39], binds to Aβ_42_ fibrils but not to tau fibrils (Fig. 1), as expected. The LCOs show different spectral characteristics when bound to tau fibrils, the detected emission peaks are for h-FTAA: 549 nm; q-FTAA: 530 nm; HS-84: 504 nm; and HS-169: 654 nm, respectively. Using the same incubation concentration and condition, h-FTAA and HS-84 give higher F.I. to tau fibrils compared to other probes (q-FTAA and HS-169).

**Fig. 1.**
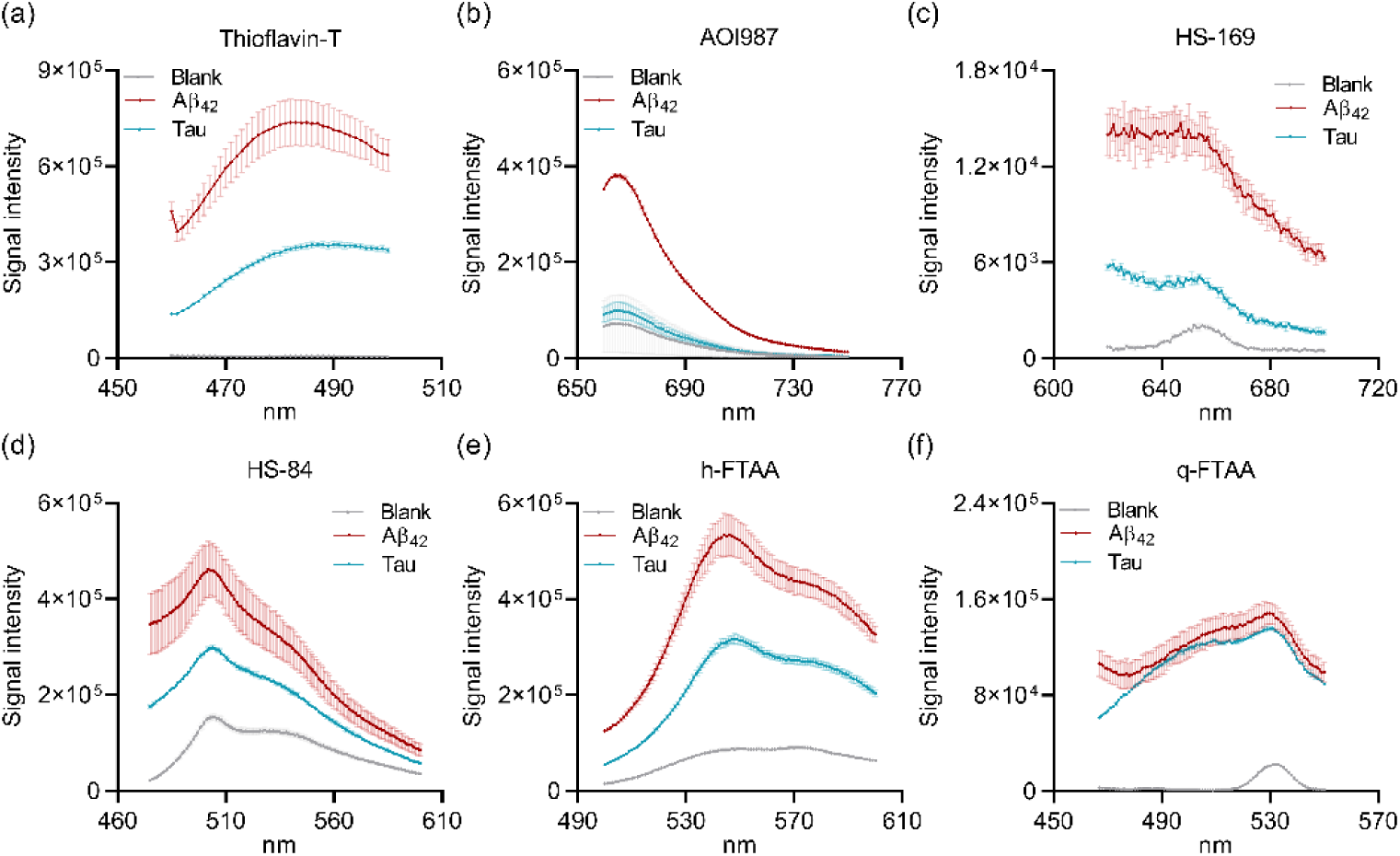
Comparison of the selectivity of different fluorescent probes to protein aggregates. Binding of (a) Thioflavin-T, (b) AOI987, and (c-f) LCOs HS-169, HS-84, h-FTAA and q-FTAA to recombinant Aβ_42_ fibril, and tau fibril.

As tauopathy deposits in human might differ in their structure and conformation from recombinant tau fibrils, we then evaluated the binding of LCOs to tauopathy on mouse brain sections of transgenic animals harboring extensive tau pathology. We choose the P301L line because these animal show an age-dependent accumulation of misfolded hyperphosphorylated tau aggregates in the brain regions (mainly hippocampus, amygdala and cortex), starting at around 7-9 months-of-age [16]. These animals do not show Aβ pathology making them an ideal tool to evaluate the LCOs selectivity towards tau [16].

Mice of 10 months-of-age displayed strong fluorescence signals in the hippocampus and cortex of the P301L mouse when brain sections were incubated with h-FTAA and HS-84 was observed (Figs. 2b, c, f, g). In comparison, weaker fluorescence signals were detected in sections stained with HS-169 (Figs. 2e) as shown previously [27], and with q-FTAA (Figs. 2a). Lambda spectrum mapping using confocal microscopy provides full spectral information from stained brain tissue slices across a range of wavelengths. Results from lambda mapping shows that the major peaks of h-FTAA and HS-84 on P301L brain tau deposits are at 543 nm and 548 nm, respectively (Figs. 2d, h). This demonstrates that h-FTAA has a similar spectral characteristic when bound to tau deposits in brain sections and recombinant tau fibrils, while HS-84 shows a blueshift in recombinant tau fibrils compared to tau deposits binding on brain sections. Taken together the results obtained on recombinant tau fibrils and P301L mouse brain, among all LCOs tested h-FTAA showed *in vitro* binding to tau fibrils and binding characteristics in brain sections of P301L. h-FTAA showed strong fluorescence emission upon binding in both, which favors its usage for *in vivo* fluorescence imaging in P301L mice.

**Fig. 2.**
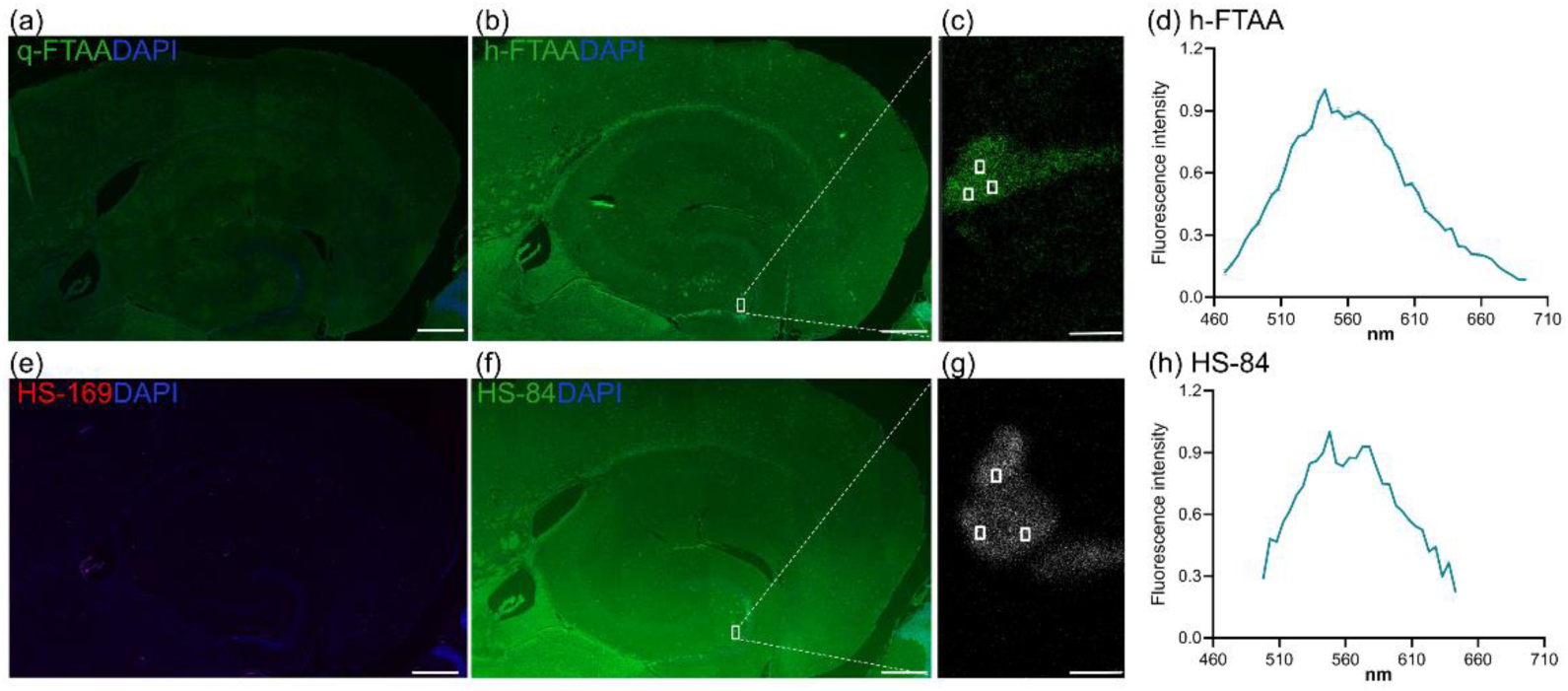
Evaluation of the cellular binding of luminescent conjugated oligothiophenes on P301L mouse brain tissue sections. Horizontal cortical brain sections from 10 months-of-age P301L mouse stained with (a) q-FTAA (green)/DAPI (blue); (b, c) h-FTAA (green)/DAPI (blue) and zoom-in; (e) HS-169 (red)/DAPI (blue), and (f, g) HS-84 (green)/DAPI (blue) and zoom-in;. Scale bar = 200 μm (a, b, e, f), 10 μm (c, g); (d, h) Lambda spectrum mapping of h-FTAA (c) and HS-84 (g) staining on P301L mouse brain slices showing the peak of binding.

### 3.2 In vivo LMI imaging of P301L and non-transgenic littermate mice

The bio-distribution and temporal dynamics of h-FTAA in mouse brain is not known in living tauopathy mouse models. Next we perform non-invasive imaging in P301L and non-transgenic littermate mice of 4-, 10- and 17-months-of-age. Mice were scanned *in vivo* by LMI and widefield imaging before and after intravenous administration of h-FTAA until 120 minutes post-injection. The h-FTAA uptake signals increased over time over the cerebrum in all groups, indicating that the probe reached the brain (Fig. 3a-c). F.I. was higher over time in the cortical region of P301L compared to non-transgenic littermate mice (p = 0.0257, two-way ANOVA, Fig. 3d). Higher F.I. (during 60-120 min) was observed in P301L compared to non-transgenic littermates (F.I. post/pre: 4.6 *vs* 1.9, *t-test*, p < 0.0001), indicating retention of probe *in vivo* in mice with tauopathy.

**Fig. 3.**
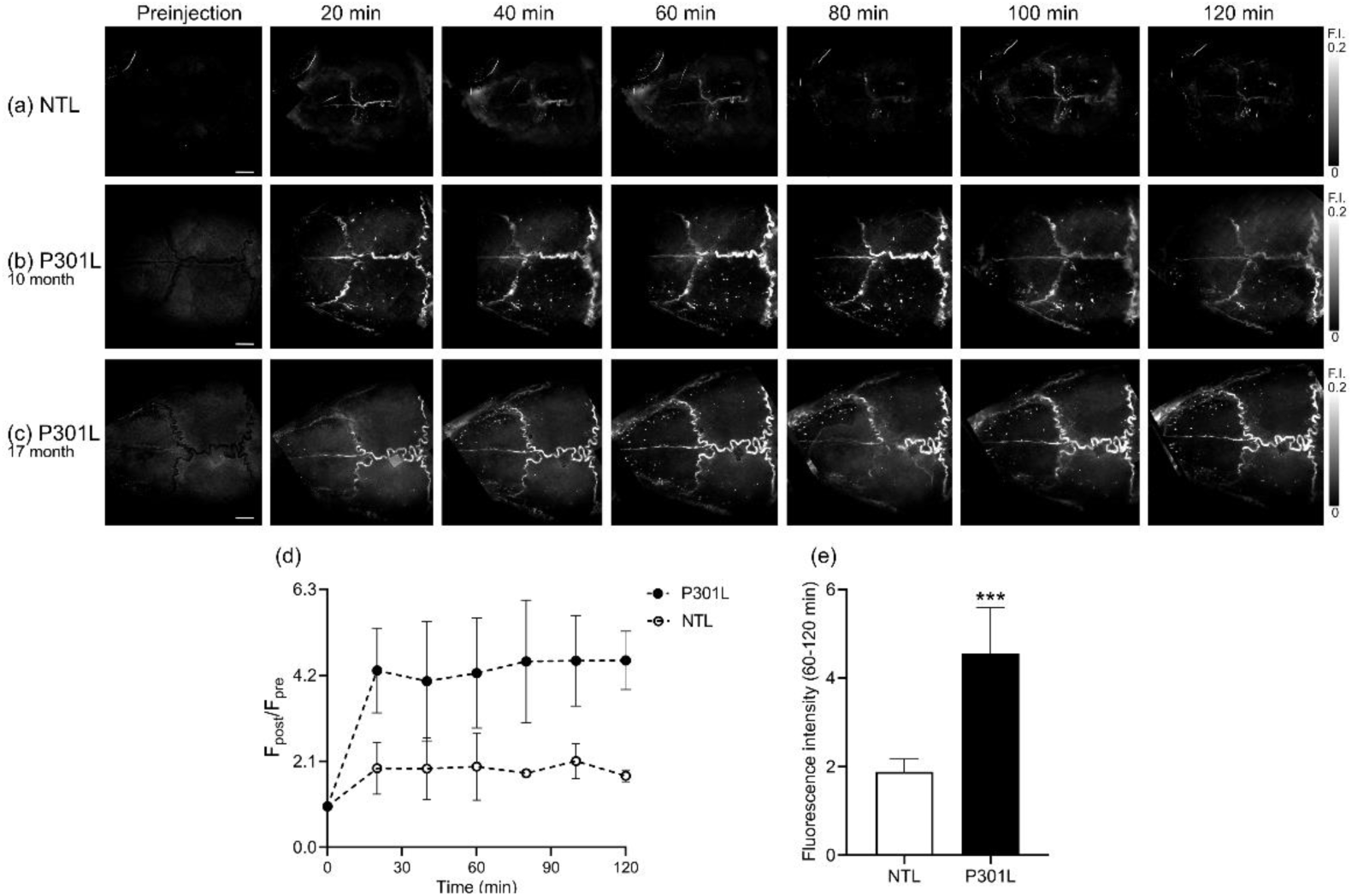
*In vivo* large-field multifocal illumination (LMI) imaging of tauopathy. (a-c) *In vivo* LMI n the brain of P301L mice at 10- and 17 months-of-age, and non-transgenic littermate (NTL) at pre-injection and following *i.v.* tail vein bolus injection of h-FTAA till120 min post-injection. Fluorescence intensity (F.I.) normalized to pre-injection values; (d) Quantification of F.I. post/pre) injection over time; NTL, n = 3, P301L n = 4; (e) Higher F.I. (60-120 min) in P301L compare to NTL mice, *** p < 0.001 t test. Scale bar = 200 μm (a-c).

### 3.3 Ex vivo LMI imaging of P301L and non-transgenic littermate mice

To confirm the *in vivo* LMI imaging finding, *ex vivo* LMI and widefield fluorescence imaging were performed on dissected whole mouse brains and on coronal brain sections at approximately 130 minutes post-injection using the same set-sup. CNR is 18 times higher using LMI imaging (Fig. 4b) compared to widefield imaging (Fig. 4a) in the *ex vivo* mouse cerebral cortex (596 *vs* 163, p < 0.0001) (Fig. 4m). The *ex vivo* LMI imaging revealed fluorescent spots in the cerebral cortex (Figs. 4b, c, g, especially the pyramidal layer Fig. 4d), hippocampus (Fig. 4h) and fluorescent structures in the skull in P301L mice. In non-transgenic littermates no such fluorescent structures were observed after h-FTAA injection. This suggests that h-FTAA binds to tauopathy deposits *in vivo* after intravenous administration (Fig. 4i-k), while it is cleared form the normal brain over the imaging period. We quantified signal intensities in different tissue compartments. A four-fold higher levels of F.I. were observed *in vivo* (p = 0.0047, *t* test, Figs, 4e, i) and *ex vivo* (p = 0.0077, Figs. 4g, k) in the cerebral cortex in the P301L compared to non-transgenic littermate mice, as well as in the *ex vivo* skull (p = 0.0108, *t* test, Figs. 4f, j). The F.I. in the *ex vivo* mouse brain and skull were approximately 80 % and 57 % respectively of that in *in vivo* mouse head (Fig. 4n).

**Fig. 4.**
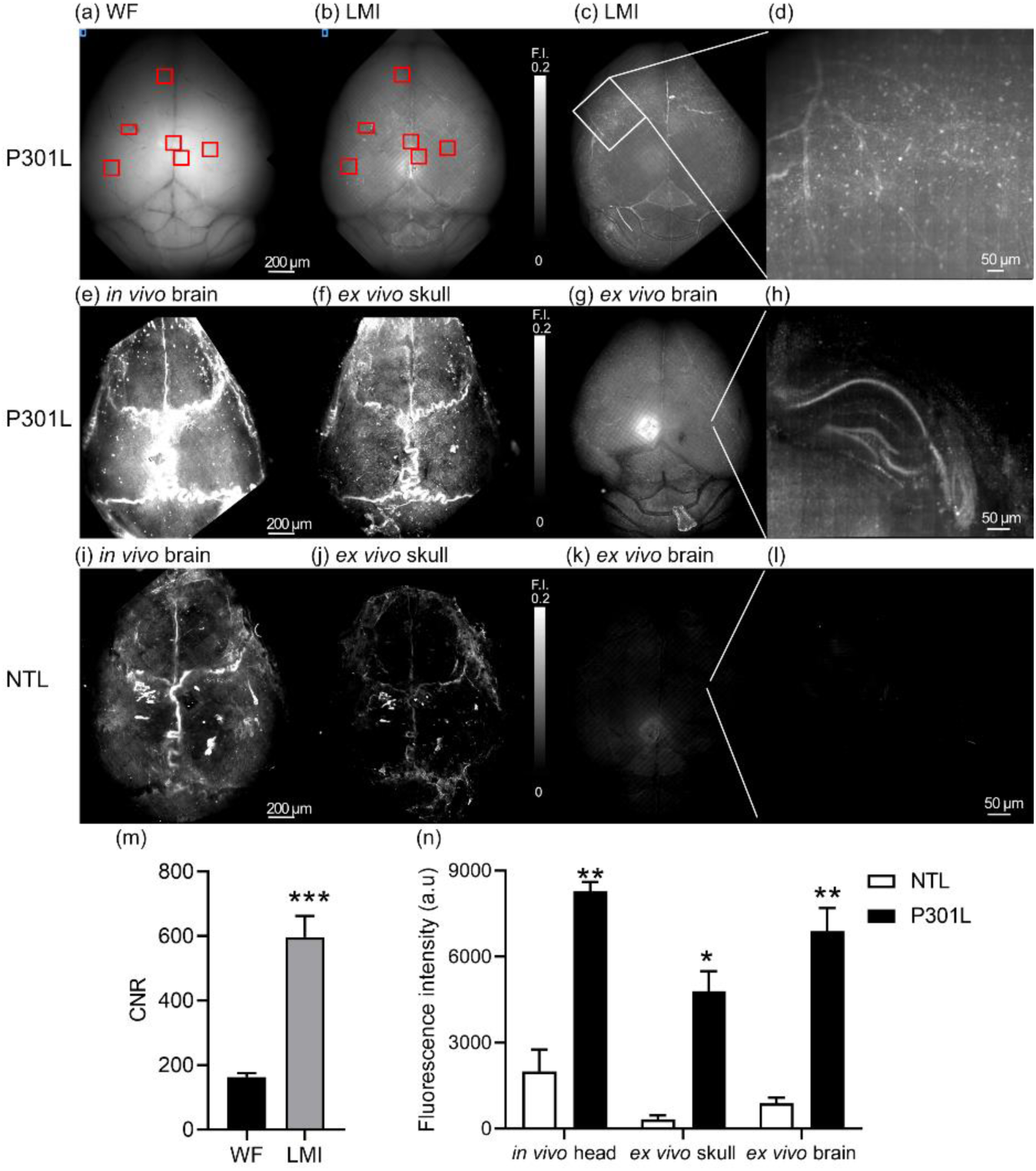
*Ex vivo* large-field multifocal illumination (LMI) imaging compared to widefield (WF) fluorescence microscopy of tauopathy in P301L mouse brain. (a-b) *Ex vivo* WF and LMI imaging of one P301L mouse brain (signal ROI (red square), and background region (blue square)); (c-d) *Ex vivo* LMI and zoom-in showing signal spots in the cortex of P301L mouse brain; (e-h) Normalized representative LMI imaging *in vivo* mouse head at 120 min post-injection, in skull, *ex vivo* whole brain and zoomed-in coronal brain slices 10 months-old P301L and (i-l) age-matched non-transgenic littermate mouse; (m) Higher contrast-to-noise ratio (CNR) in LMI imaging compared to widefield imaging analyzed using over P301l mouse *ex vivo* cortex; (n) Quantification of signals in the brain and in the skull in P301L and non-transgenic littermate mice; * *p* < 0.05, ** *p* < 0.01, *** *p* < 0.001 *t* test (m, n).

### 3.4 Specificity of h-FTAA for detection of tauopathy on brain sections from P301L mice

To validate the *in vivo* finding and to assess the specificity of h-FTAA binding to tau deposits in mouse brains, we performed immunohistochemistry on horizontal brain tissue sections from P301L mice and non-transgenic littermates using h-FTAA with phosphorylated tau AT-8 and AT-100 antibodies (counterstained with DAPI). P301L mice have only tauopathy deposits, and no Aβ aggregates (which is also positive to h-FTAA) [16, 38, 40]. In the P301L mouse brain, h-FTAA co-localized with AT-8 positive cells, which binds to sarkosyl-insoluble tau and soluble hyperphosphorylated tau [41] (Fig. 5a, b). In addition, h-FTAA co-labeled neuronal inclusion that were positive for AT-100, which only recognizes sarkosyl-insoluble, but not soluble tau (Figs. 6c) [41]. Distribution in the cortex and hippocampus especially CA1 pyramidal neuron, cell bodies, apical dendrites, and in mossy fiber projection, moderate in the dentate gyrus granule cells were observed. No signal was observed on h-FTAA/AT-8 (Figs. 6d, e) and h-FTAA/AT-100 (Figs. 6f, g) stained non-transgenic littermate mouse brain sections, indicating specific targeting of the probe to tau deposits.

**Fig. 5.**
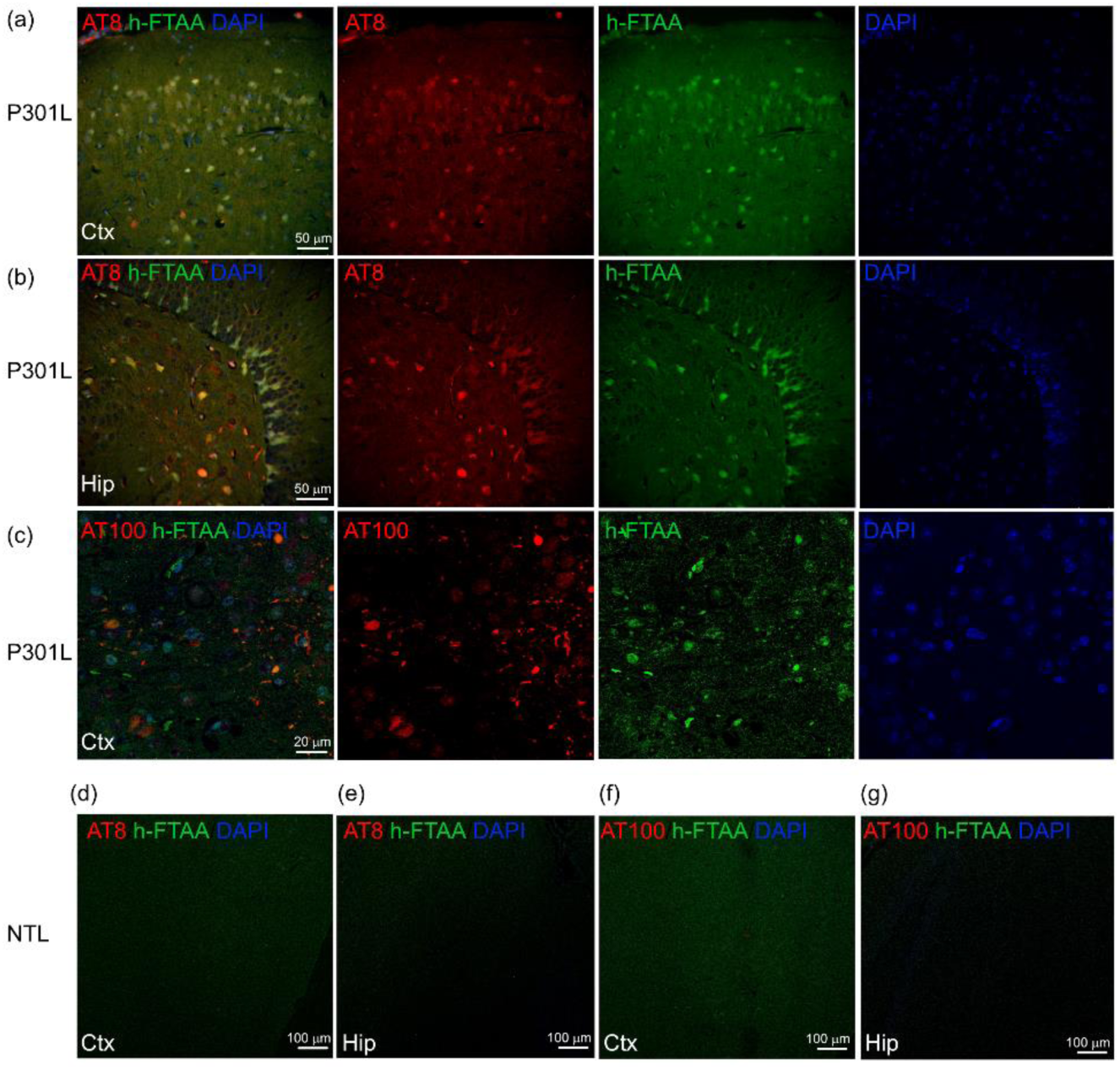
Specificity of h-FTAA for tauopathy in P301L mouse brain tissue sections. (a-c) Immunohistochemistry of horizontal brain sections from the cortex and hippocampus of P301L mouse, demonstrating co-localization of AT-8, AT-100 and h-FTAA to tauopathy in P301L mouse brain; (d-g) No specific signals in the cortex and hippocampus of non-transgenic littermate mouse brain are stained by AT-8/ h-FTAA or AT-100/h-FTAA. (a,b,d,e) AT-8 (red), (c, f, g) AT-100 (red), h-FTAA (green), DAPI (blue).

## 4. Discussion

Developing tools for non-invasive detection of tau deposits at high-resolution in animal models of tauopathy is critical for understanding disease mechanism [42] and for translational development of tau-targeted therapies and diagnostics [6, 43–45]. Here we demonstrated a novel *in vivo* transcranial LMI imaging approach to map tau deposits at the whole brain scale in P301L mouse model of FTLD tauopathy using h-FTAA that binds with high sensitivity and specificity to tau aggregates.

Thanks to the inherent advantage of laser beam scanning, the proposed LMI imaging method can effectively reject the out-of-focus light, enabling minimally invasive imaging *in vivo* without employing craniotomy, an essential advantage especially when it comes to longitudinal study of disease progression, such as tauopathy spreading mechanism in the brain [46]. Due to improvement in the spatial resolution of image scanning microscopy, the LMI imaging approach can achieve 6 μm resolution across the entire mouse cortex. Compared to laser scanning confocal microscopy, LMI imaging is a highly parallelized technique employing hundreds of illumination foci, thus enabling fast imaging, which is crucial for mitigating motion artifacts in *in vivo* studies. In addition, the LMI imaging approach is optimally suited for imaging large objects up to a centimeter scale, which is not attainable with multi-photon microscopy methods that are further hindered by the lack of optimal labels with large absorption cross section and longer absorption/emission wavelengths. Thus, the set-up is ideally suited to detect immunological and vascular events in the mouse brain using dedicated probes [47–49].

Tau imaging has been challenging, due to the structural diversity of tau isoforms, the lack of specificity and off-target binding property of many tau imaging probes [50]. Recently a new family of LCOs were described that detect aggregated proteins such as Aβ and tau fibrils [32]. The LCOs h-FTAA [51], and HS-84 [27] have been used to detect tauopathy in postmortem human brain tissue and in intravital brain imaging of tau deposits in P301L mouse models using two-photon microscopy with a cranial window. Here, we demonstrated the *in vivo* transcranial imaging of taupathy in a non-invasive manner.

We first compared four LCOs for their suitability for *in vivo* LMI imaging. All probes have different spectral characteristics, where the emission peak of HS-169 would render the probe most suitable for deep tissue fluorescence imaging with less absorption and scattering compared to other probes with lower emission wavelengths. However, q-FTAA and HS-169 showed low fluorescence upon binding to tau aggregates in *in vitro* binding assay, while h-FTAA and HS-84 showed bright fluorescence. Moreover, we found that all LCOs did not only bind to tau but also to Aβ42 fibrils. They bind to protein aggregates having an extensive cross β-pleated sheet conformation [31], as many other tau probes such as PBB3, Thioflavin S, Congo Red, and its derivative FSB [30, 52–54]. However, compared to dyes such as Thioflavin S and Congo Red, LCOs have a stronger fluorescence emission. The lack of selectivity restricts the use of LCOs in animal models of mixed Aβ and tau pathologies. It has been shown that some LCOs undergo a structural restriction upon protein binding and a shift of the emitted light occurs, dependent on the target protein and its conformation, which would allow discriminating protein deposits based on their spectroscopic signatures *ex vivo* [32].

We evaluated the binding of LCOs to tauopathy on mouse brain sections of P301L mice. Binding of probes to tau fibrils might be different to probe binding in brain tissue due to the structural diversity of tau aggregates. For example, it was reported recently that HS-169 binds to Aβ and tau fibrils [30], but does not recognize tau deposits in the brain of P301L mice (CaMKII) [27]. We found low fluorescence signals of q-FTAA and HS-169 on brain sections from P301L mice, while sections subjected to HS-84 and h-FTAA had intense and localized fluorescence. Since the lambda spectrum mapping showed a blueshift of HS-84 in tau binding on brain sections compared to h-FTAA, we opted for the use of h-FTAA for *in vivo* LMI imaging.

We investigated whether the h-FTAA would enter the brain across the blood-brain barrier in amounts sufficient for *in vivo* brain imaging using LMI. An increase in fluorescence was observed after intravenous administration of probe and uptake of probe to the brain of P301L mice was apparent after excising the brain from the skull. Moreover, we used the high-resolution capability of the LMI system and detected fluorescent circular structures in the excised brain. Tauopathy deposits in P301L (Thy1.2) mice start in entorhinal-hippocampal regions from approximately 7 months-of-age [16], are most pronounced in the cortex, amygdala and hippocampus, moderate in brain stem and striatum, and negligible in the cerebellum. The cortical signals detected by LMI imaging *in vivo* and *ex vivo* using h-FTAA are in accordance with immunohistochemical staining results (Figs. 4-6), and with the known tau distribution in the P301L mouse brain [40]. We observed that part of the signals originated for the skull in the tauopathy mouse brain. Due to the wavelength of the probe, hemoglobin may also contribute to the signal.

The skull impairs high-resolution optical imaging of the mouse cortex *in vivo* due to its strong scattering of the light. Using non-invasive fluorescence imaging set up, somatosensory-evoked rapid calcium transients in the GCaMP6f brain (peak excitation: 488 nm, emission: 512 nm) *in vivo* was detected despite blurred fluorescence images [55]. In the proposed LMI imaging method, the scattering of the excitation light in the surrounding tissue was mitigated by introducing the multifocal illumination. The scattering of the emitted fluorescence signal was reduced during image reconstruction with only the local maxima in the image were extracted and further superimposed in the final reconstructed image. Chronic large cranial window techniques and skull optical clearing solutions were proposed to reduce the skull scattering coefficient [56]. In addition, detection of tau deposits from the brain could be improved by using red shifted tau imaging probes with longer excitation and emission wavelengths. Further studies are needed to clarify the signals within skull in this tauopathy mouse model.

We screened LCOs and assessed the specificity of h-FTAA as a tau probe and whether h-FTAA can detect pathological tau in all tauopathies in brain tissue slices from P301L mouse using immunohistochemistry. We used antibodies that recognize insoluble (AT-100), and filamentous and soluble (AT-8) forms of hyperphosphorylated tau. The h-FTAA localized well both with AT-8 and AT-100 positive cells. In comparison, no probe accumulation was observed in the brain of non-transgenic littermates that were tau negative, suggesting a specificity of the probe *in vivo*. We observed that h-FTAA were not washed away again with time, potentially suitable for longitudinal imaging of tauopathy deposits *in vivo* [27]. HS-84 that showed a blue shift probably due to selectivity for *in vitro* tau fibrils which have an entirely different structure than *in vivo* murine tau fibrils as demonstrated by the cryogenic electron microscopy structures [57].

In conclusion, we demonstrated high-resolution whole brain *in vivo* transcranial LMI fluorescence imaging for tauopathy in P301L mouse model of FTLD using h-FTAA. Such platform will provide insight in mechanistic studies of tau spreading [58, 59] and clearance [60], as well as in longitudinal monitoring of tau targeting therapeutics [61] in tauopathy animal models.

